# Mycobacterial infection-induced miR-206 inhibits protective neutrophil recruitment via the CXCL12/CXCR4 signalling axis

**DOI:** 10.1101/2020.12.14.422665

**Authors:** Kathryn Wright, Kumudika de Silva, Karren M. Plain, Auriol C. Purdie, Tamika A Blair, Iain G Duggin, Warwick J. Britton, Stefan H. Oehlers

## Abstract

Pathogenic mycobacteria actively dysregulate protective host immune signalling pathways during infection to drive the formation of permissive granuloma microenvironments. Dynamic regulation of host microRNA (miRNA) expression is a conserved feature of mycobacterial infections across host-pathogen pairings. Here we examine the role of miR-206 in the zebrafish model of *Mycobacterium marinum* infection, which allows investigation of the early stages of granuloma formation. We find miR-206 is upregulated following infection by pathogenic *M. marinum* and that antagomir-mediated knockdown of miR-206 is protective against infection. We observed striking upregulation of *cxcl12a* and *cxcr4b* in infected miR-206 knockdown zebrafish embryos and live imaging revealed enhanced recruitment of neutrophils to sites of infection. We used Crispr/Cas9-mediated knockdown of *cxcl12a* and *cxcr4b* expression and AMD3100 inhibition of Cxcr4 to show that the enhanced neutrophil response and reduced bacterial burden caused by miR-206 knockdown was dependent on the Cxcl12/Cxcr4 signalling axis. Together, our data illustrate a pathway through which pathogenic mycobacteria induce host miR-206 expression to suppress Cxcl12/Cxcr4 signalling and prevent protective neutrophil recruitment to granulomas.

**Author summary:** Mycobacterial infections cause significant disease burden to humans and animals, the most widely known example being tuberculosis which has killed more humans than any other infectious disease throughout history. Infectious mycobacteria are highly evolved to hijack host processes, including the very immune cells tasked with destroying them. microRNAs are host molecules that control wide-ranging programs of host gene expression and are important in the immune response to infections. Here we use the zebrafish model of mycobacterial infection to determine the role of the infection-induced microRNA miR-206 in the host response to infection. We found pathogenic mycobacteria trigger the host to produce more miR-206 in order to suppress the otherwise protective recruitment of neutrophils to sites of infection via the host Cxcl12/Cxcr4 signalling pathway. Our study provides new insight into the role of mycobacterial infection-induced miR-206 function in the context of a whole host.

## Introduction

Pathogenic mycobacteria, including the causative agents of tuberculosis and leprosy, are capable of appropriating host signalling and immune pathways to increase their survival and establish chronic infection in cell-rich granulomas, which support mycobacterial growth and latent survival (1, 2).

microRNA (miRNA) are short, non-coding RNA of approximately 22 nucleotides that can post-transcriptionally regulate gene expression and transcript abundance through “gene silencing”. miRNA bind to the untranslated region (UTR) of mRNA to regulate the stability of target genes through degradation or suppression, reducing protein translation (3). Expression of miRNA is dynamically regulated in mycobacterial infection, suggesting a key role in the host response to infection by the modulation of downstream genes and protein expression (4–6).

miR-206 is a member of the muscle-associated myomiR family and is characteristically associated with myoblast differentiation and muscle development (7-10). However, miR-206 has been recently found to be differentially regulated in mycobacterial infection of THP-1 leukocytic cells (11). Infection of THP-1 cells with *Mycobacterium tuberculosis* revealed a role for miR-206 in the regulation of proinflammatory cytokine responses through reducing TIMP3 expression (11). miR-206 has also been implicated in viral pathogenesis, reducing replication of influenza virus, and in neuroinflammation (12, 13). While miR-206 clearly has a diverse range of biological functions, the *in vivo* role of miR-206 during mycobacterial infection remains undetermined.

Here we use the zebrafish-*Mycobacterium marinum* model to investigate the *in vivo* function of dre-miR-206-3p (miR-206) in mycobacterial infection, and the impact on host gene expression and neutrophil recruitment. Zebrafish are an established model for the investigation of host-mycobacteria interactions, and to evaluate host gene function, including the role of chemokine signalling (14-16). Regulation of neutrophil based inflammation and motility by miRNA has also been investigated using a zebrafish model (17-21), highlighting the applicability of the zebrafish model to study conserved miRNA functions within host-pathogen interactions.

## Results

### Zebrafish miR-206 expression is responsive to *M. marinum* infection

To determine if miR-206 is responsive to *M. marinum* infection in zebrafish, embryos were infected with *M. marinum* via caudal vein injection and miR-206 expression was measured by quantitative (q)PCR at 1, 3, and 5 days post infection (dpi). Infection with *M. marinum* increased miR-206 expression at 1 and 3 dpi, but decreased miR-206 expression at 5 dpi compared to uninfected controls (Fig 1A).

**Fig 1.**
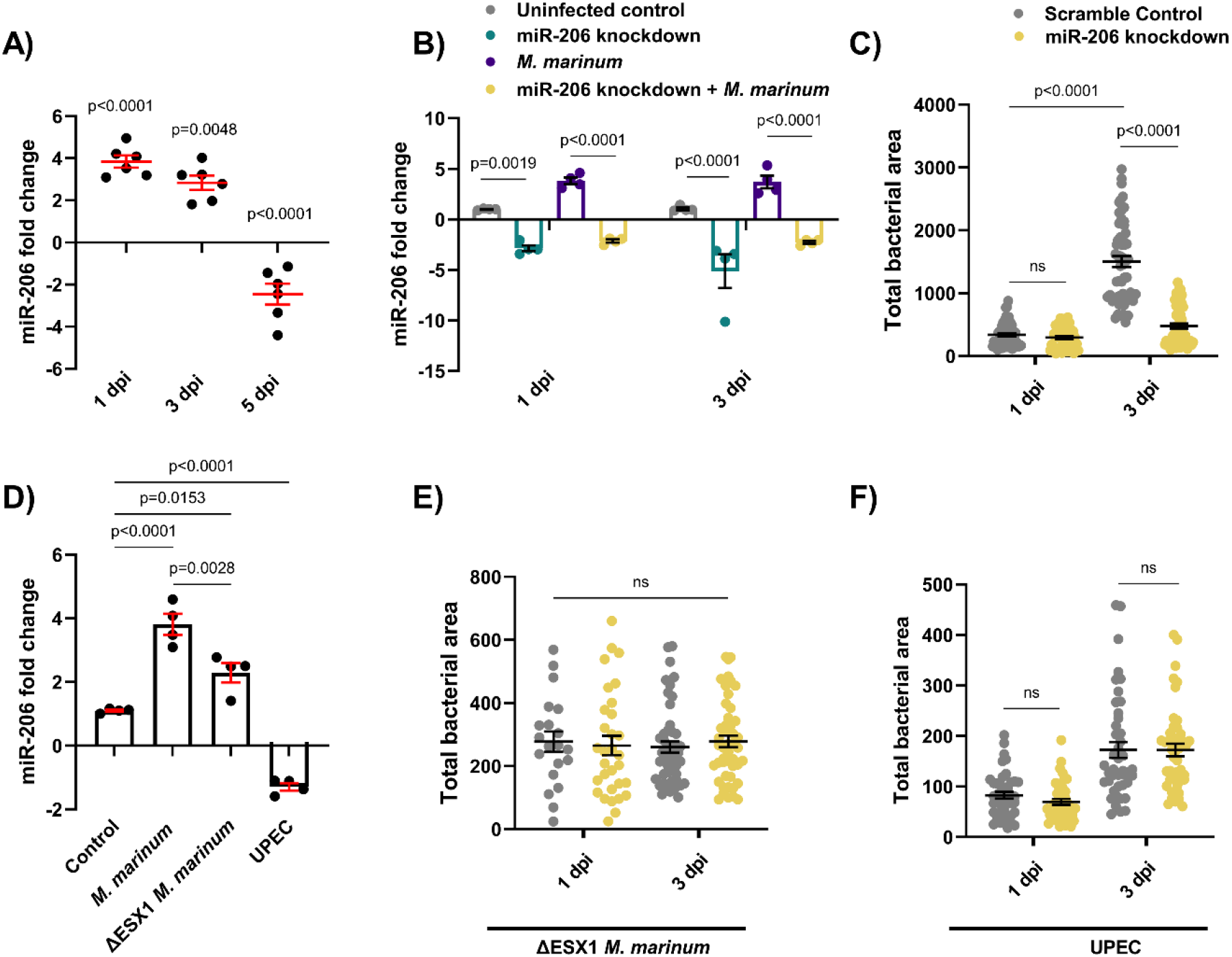
Infection-induced miR-206 expression alters bacterial burden. (A) Expression of miR-206 analysed by qPCR at 1, 3, and 5 dpi. (B) Expression of miR-206 in uninfected and infected antagomir-injected embryos (miR-206 knockdown). (C) *M. marinum* burden in miR-206 knockdown embryos at 1 and 3 dpi. (D) Expression of miR-206 at 1 dpi following infection with either wild-type (WT) *M. marinum*, ΔESX1 *M. marinum*, or UPEC. (E) ΔESX1 *M. marinum* burden in miR-206 knockdown embryos at 1 and 3 dpi. (F) UPEC burden in antagomir-injected embryos at 6 hpi and 1 dpi. Each data point represents a single measurement, with the mean and SEM shown. For qPCR analysis, each data point represents 10 embryos, and contains 2 biological replicates. Bacterial burden analysis data points (WT *M. marinum*, ΔESX1 *M. marinum*, and UPEC) represent individual embryos (n=40-50 embryos per group) and are representative of 2 biological replicates.

### AntagomiR abrogates infection-induced miR-206 expression

To determine the efficacy of antagomiR-mediated miRNA knockdown during *M. marinum* infection, embryos were injected with miR-206 antagomiR at the single-cell stage and infected at 1.5 days post fertilisation (dpf). miR-206 expression levels were analysed at both 1- and 3-days post infection (dpi) and were increased in *M. marinum* infected embryos compared to control uninfected (p=0.027 and p=0.039 respectively). AntagomiR knockdown effectively reduced the miR-206 level relative to infected embryos at both timepoints, demonstrating effective knockdown of infection-induced miR-206 expression by antagomiRs (Fig 1B).

### miR-206 knockdown reduces *M. marinum* burden

As miR-206 was modulated during infection, its effect on disease was assessed through assessing the bacterial burden following antagomiR knockdown. There was no difference in bacterial burdens between miR-206 knockdown and control embryos at 1 dpi, however by 3 dpi, knockdown embryos had a significantly lower burden than control embryos. (Fig 1C).

### Infection-induced miR-206 upregulation is driven by mycobacterial virulence factors

To investigate whether the decreased bacterial burden in miR-206 knockdown embryos was a general response to foreign pathogens or a more directed response, embryos were infected with either ΔESX1 *M. marinum* or uropathogenic *Escherichia coli* (UPEC). ΔESX1 *M. marinum* lack the key type VII secretion system and are far less virulent as they are unable to lyse host cell membranes to escape the phagosome (22). In comparison to mycobacteria, UPEC cause an acute sepsis infection and are an example an extracellular bacterium.

Expression of miR-206 was analysed by qPCR in embryos infected with WT *M. marinum*, ΔESX1 *M. marinum*, or UPEC at 1 dpi (Fig 1D). Infection with ΔESX1 *M. marinum* increased miR-206 expression, however this response was less than the level induced by infection with virulent WT *M. marinum*. Conversely, miR-206 was decreased in embryos infected with UPEC.

ΔESX1 *M. marinum* infection burdens were unaffected by miR-206 knockdown at either 1 or 3 dpi (Fig 1E). Similarly, there was also no difference in UPEC burden levels in miR-206 knockdown embryos at either 6 hours post infection (hpi) or 1 dpi despite an increase in infection between timepoints (Fig 1F). These results indicate that the impact on bacterial burden in miR-206 knockdown embryos is driven by *M. marinum* virulence factors.

### miR-206 target mRNA gene expression patterns are conserved during *M. marinum* infection of zebrafish

To further investigate the functional relevance of miR-206 in mycobacterial infection, a list of potential mRNA target genes was compiled through published experimentally observed targets and bioinformatic target prediction algorithms (23-26).

Expression of selected potential target genes of miR-206 was analysed by qPCR at 2 dpf, with increased expression in knockdown samples expected to indicate targeting by miR-206 (Fig 2A). From the expression profiling data, *cxcl12a* and *cxcr4b* were considered to be likely targets of miR-206. Expression of these genes was increased by *M. marinum* infection and in both knockdown treatments, suggesting they may be active during infection and contributing to the decreased bacterial burden observed in miR-206 knockdown embryos. These genes were also of particular interest as the Cxcl12/Cxcr4 pathway has been previously implicated in zebrafish immunity (14, 15). Expression of *cxcr4a, cxcr4b, cxcl12a* was analysed at 3 dpi, showing that knockdown of miR-206 significantly increased the transcript abundance of *cxcr4b* and *cxcl12a* in infected embryos compared to *M. marinum* infection alone and uninfected controls (Fig 2B-D).

**Fig 2.**
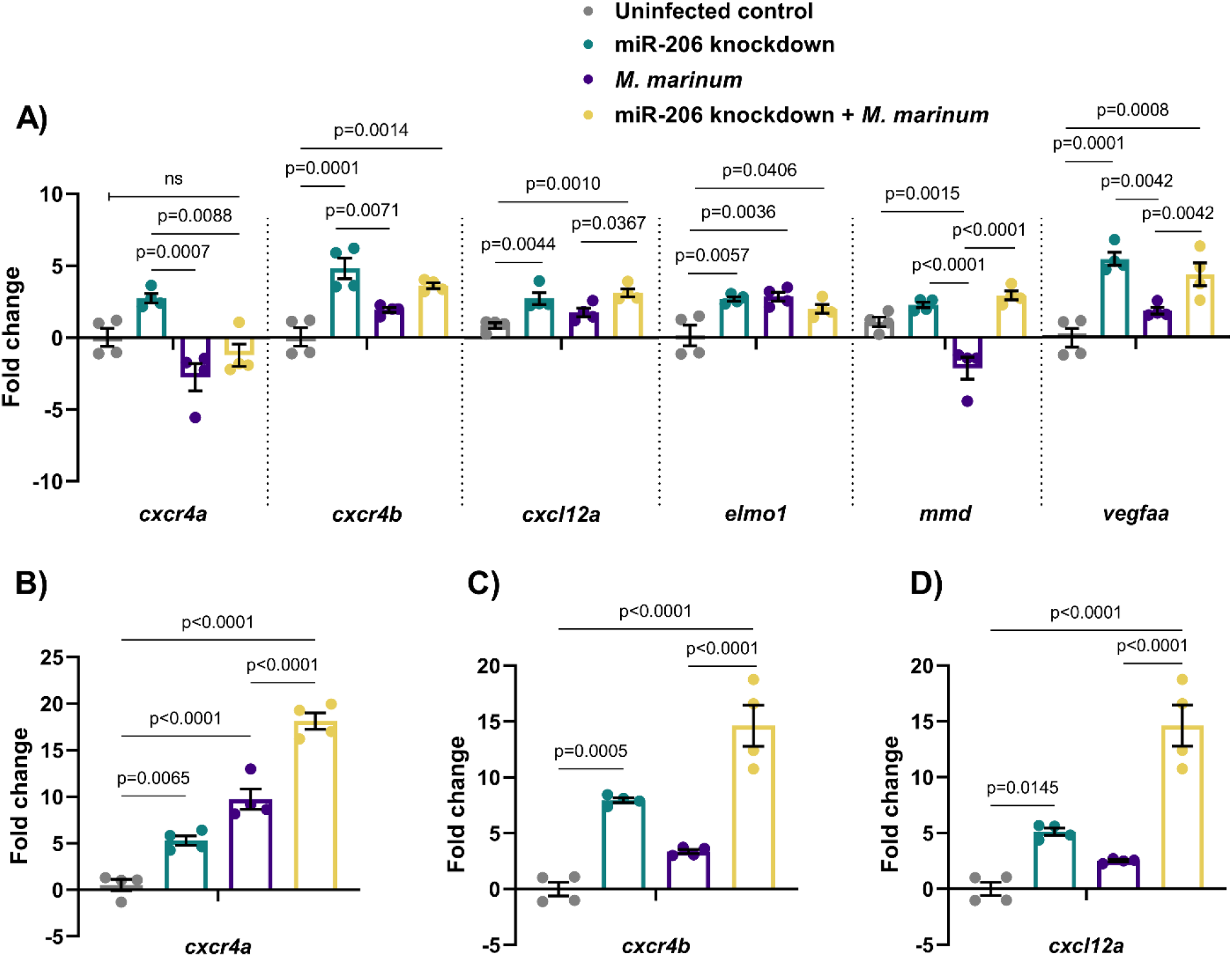
Expression profiles of potential mRNA targets of miR-206. (A) Expression of candidate target genes measured by qPCR at 1 dpi in miR-206 knockdown embryos. (B-D) Expression of zebrafish CXCL12/CXCR4 pathway ortholog genes at 3 dpi. Each data point represents a single measurement of 10 pooled embryos and 2 biological replicates, with the mean and SEM shown.

### Knockdown of miR-206 increases neutrophil response to *M. marinum* infection

As miR-206 knockdown increased the expression of *cxcl12a* and *cxcr4* genes, which are involved in neutrophil migration and retention of cells at sites of infection and inflammation (19, 27-29), the neutrophil response to *M. marinum* infection in miR-206 knockdown treated embryos was assessed by live imaging of transgenic *Tg(lyzC:GFP)^nz117^* or *Tg(lyzC:DsRed2)^nz50^* embryos, where neutrophils are fluorescently labelled.

First, static imaging was performed at 1 and 3 dpi to measure total neutrophil numbers in infected embryos. At each timepoint, miR-206 knockdown embryos had a significantly higher total number of neutrophils compared to control (Fig 3A-B). We noted strong overlap of neutrophils with *M. marinum* but the location of granulomas around the caudal haematopoietic tissue confounded quantification of granuloma-associated neutrophils.

**Fig 3.**
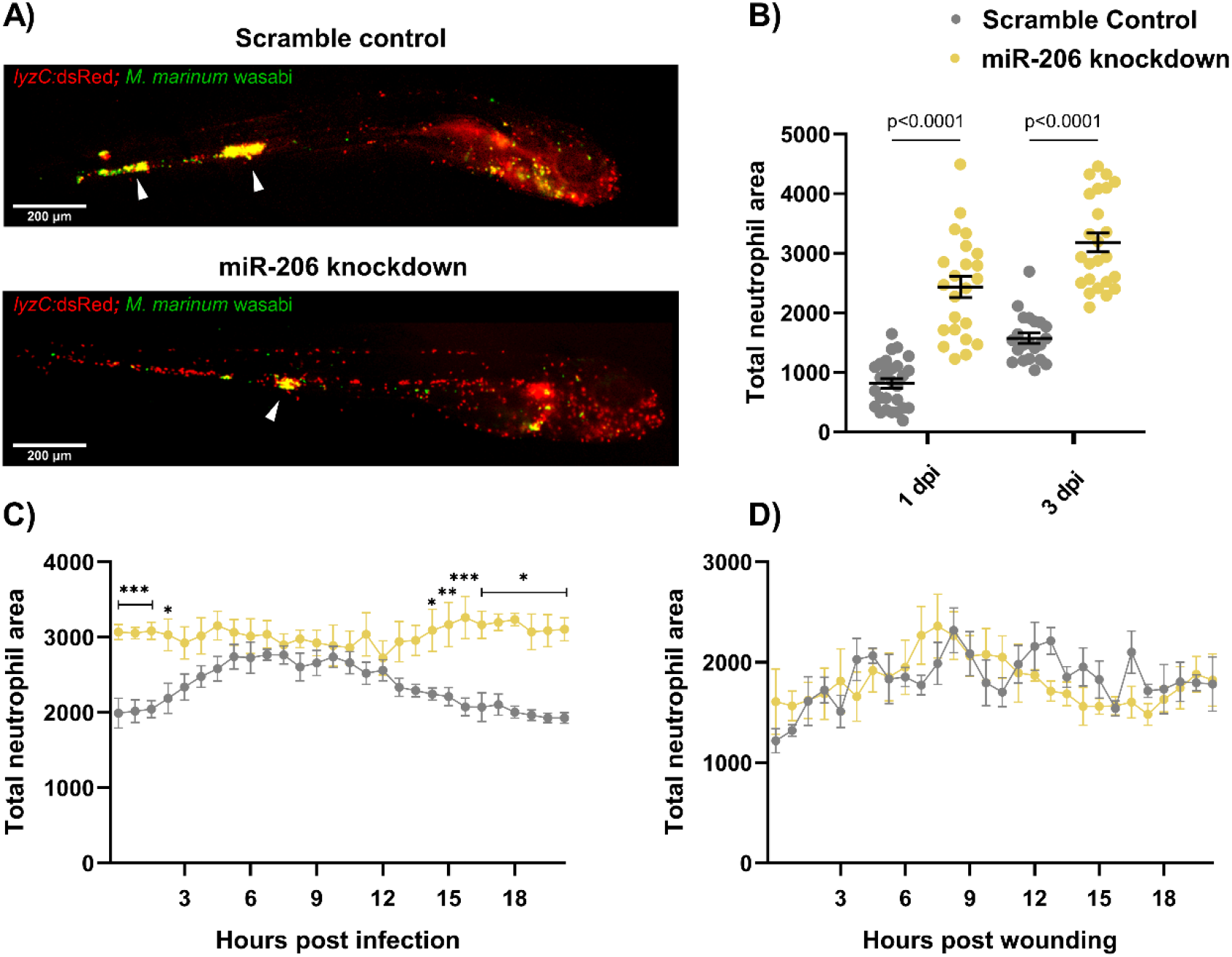
Infection-induced miR-206 expression alters the host neutrophil response. (A) Representative images of infection phenotype at 3 dpi in control and miR-206 knockdown embryos. White arrows indicate bacterial foci. Neutrophils are red (*lyzC:dsred*) and *M. marinum* is green (wasabi); co-localisation is indicated by yellow fluorescence. (B) Measurement of whole-body neutrophil fluorescent area at 1 and 3 dpi in miR-206 knockdown embryos. (C) Measurement of neutrophil levels following trunk infection with *M. marinum* in miR-206 knockdown embryos. (D) Measurement of neutrophil recruitment to a sterile tail fin wound in miR-206 knockdown embryos. Each data point represents a single measurement, with the mean and SEM shown. For time-lapse imaging, each data point represents the mean of 6 foci of infection from 6 separate embryos. Bacterial burden analysis was performed on 15-20 embryos per treatment. Graphs are representative of 2 biological replicates. * P < 0.05, ** p < 0.01, *** p < 0.001

To determine if miR-206 knockdown increased the number of infection-associated neutrophils, embryos were injected with *M. marinum* into the trunk (away from the caudal haematopoietic tissue) and subjected to time-lapse imaging (30). Knockdown embryos had significantly more neutrophils at the site of infection for the first 2.5 hours of infection compared to control infected embryos (Fig 3C). While neutrophil migration in control infected embryos began to wane at approximately 12 hpi (S1 video), the response in the knockdown embryos was sustained and higher numbers of neutrophils were maintained at the site of infection (S2 video).

To examine if the increased mobilisation of neutrophils in the *M. marinum*-infected miR-206 knockdown embryos was dependent on mycobacterial infection cues or an intrinsic feature of neutrophils in miR-206-depleted animals, we assessed neutrophil migration to a sterile tail fin wound as an example of a non-infectious inflammatory stimulus (Fig 3D). The number of neutrophils at the wound site did not significantly differ between scramble control (S3 video) and miR-206 knockdown embryos (S4 video), indicating the increased neutrophil response observed in trunk infections is *M. marinum* infection-dependent.

### The Cxcr4/Cxcl12 signalling axis is downstream of miR-206 in *M. marinum* infection

To confirm the hypothesised link between the observed increased transcription of *cxcr4b* and *cxcl12a* and reduced bacterial burden through an increased neutrophil response early in infection, both genes were targeted for knockdown by Crispr-Cas9. As both *cxcr4b* and *cxcl12* are involved in neutrophil migration and haematopoiesis, a reduction in their expression was expected to result in a reduced neutrophil response to infection and therefore an increased bacterial burden, reducing the protective effect of miR-206 knockdown.

Static imaging at 3 dpi revealed that double knockdown of *cxcr4b* and miR-206 ablated the increased neutrophil number associated with miR-206 knockdown (Fig 4A). Furthermore, addition of *cxcr4b* knockdown to miR-206 knockdown dampened the miR-206 knockdown-induced increase in neutrophil recruitment to a trunk infection and increased bacterial burden back to control levels (Fig 4B-C). The effect observed in the double knockdown is consistent with a reduction in Cxcr4 and therefore the neutrophil response in infection via haematopoiesis and chemoattraction. This suggests the miR-206 associated increase in *cxcr4b* is contributing to the enhanced neutrophil migration and reduced bacterial burden.

To further confirm involvement of Cxcr4 downstream of miR-206, the CXCR4 antagonist AMD3100 was used to pharmacologically block Cxcr4 signalling. AMD3100 treatment reduced the total neutrophil numbers in all treatment groups, and as expected, whole-body neutrophil counts were reduced in miR-206 knockdown embryos that were also treated with AMD3100 compared to miR-206 knockdown alone (Fig 4D). AMD3100 treatment decreased neutrophil recruitment to the site of infection compared to both control infected and miR-206 knockdown infected embryos (Fig 4E). Bacterial burden was increased in AMD3100 treated embryos and AMD3100 treatment of miR-206 knockdown embryos restored bacterial burden to control levels (Fig 4F).

**Fig 4.**
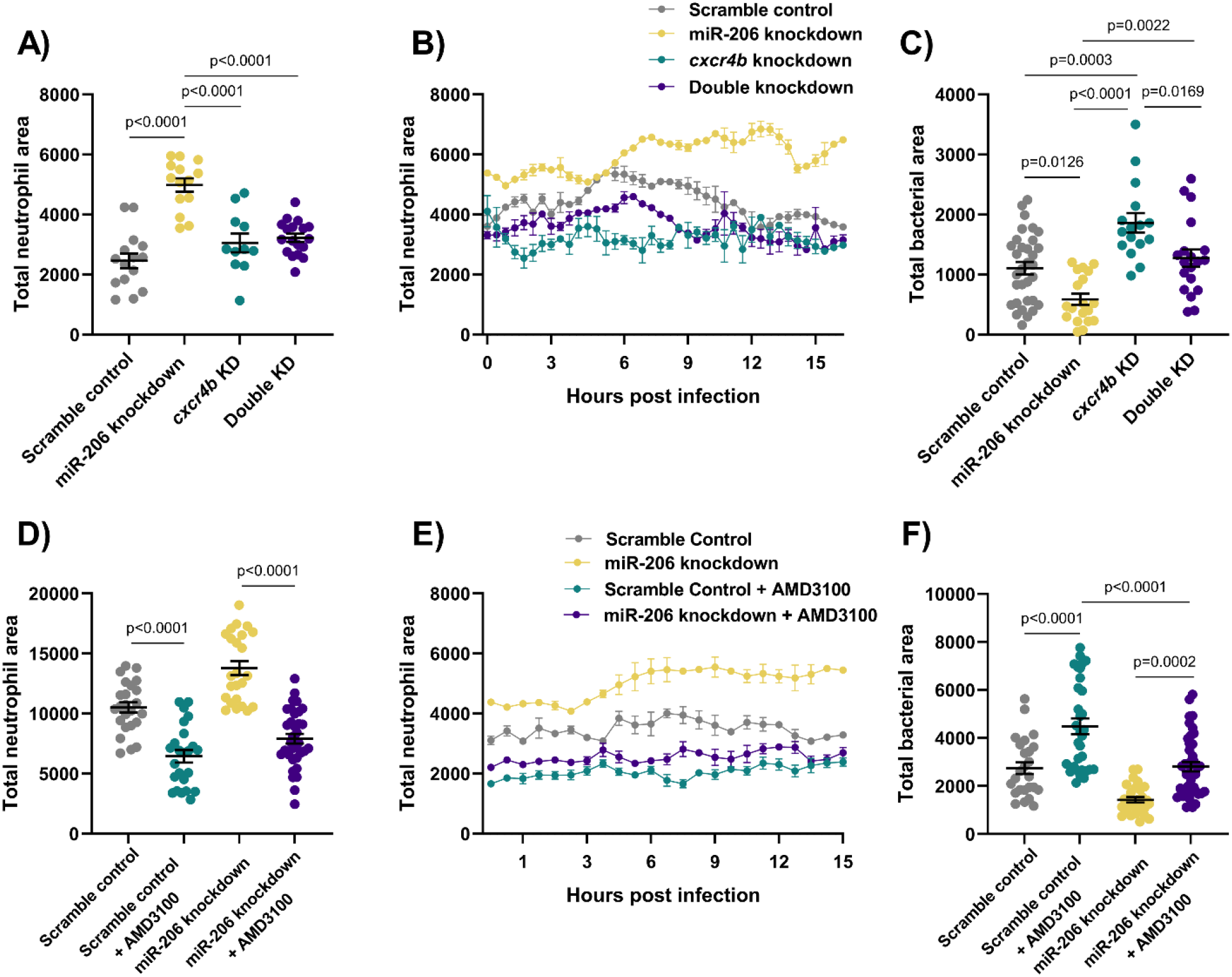
Cxcr4 reduction places the Cxcl12/Cxcr4 signalling axis downstream of miR-206. (A) Whole body neutrophil counts at 3 dpi of *cxcr4b* and double (*cxcr4b* and miR-206) knockdown embryos. (B) Measurement of neutrophil levels following trunk infection with *M. marinum* in double knockdown embryos. (C) Bacterial burden at 3 dpi in *M. marinum*-infected double knockdown embryos. (D) Whole body neutrophil counts at 3 dpi in miR-206 knockdown embryos treatment with AMD3100. (E) Measurement of neutrophil recruitment to *M. marinum* following trunk injection in miR-206 knockdown embryos treatment with AMD3100. (F) Bacterial burden at 3 dpi in miR-206 knockdown embryos treatment with AMD3100. Each data point represents a single measurement, with the mean and SEM shown. For time-lapse imaging, each data point represents the mean of 6 foci of infection from 6 separate embryos. Bacterial burden analysis was performed on 15-25 embryos per treatment. Graphs are representative of 2 biological replicates, except for AMD3100 data, which is a single biological replicate.

Consistent with the results of *cxcr4b* knockdown, addition of *cxcl12a* knockdown to miR-206 knockdown decreased the total number of neutrophils and number of neutrophils recruited to sites of infection compared to miR-206 knockdown alone (Fig 5A-B), and increased the bacterial burden compared to miR-206 knockdown alone (Fig 5C).

**Fig 5.**
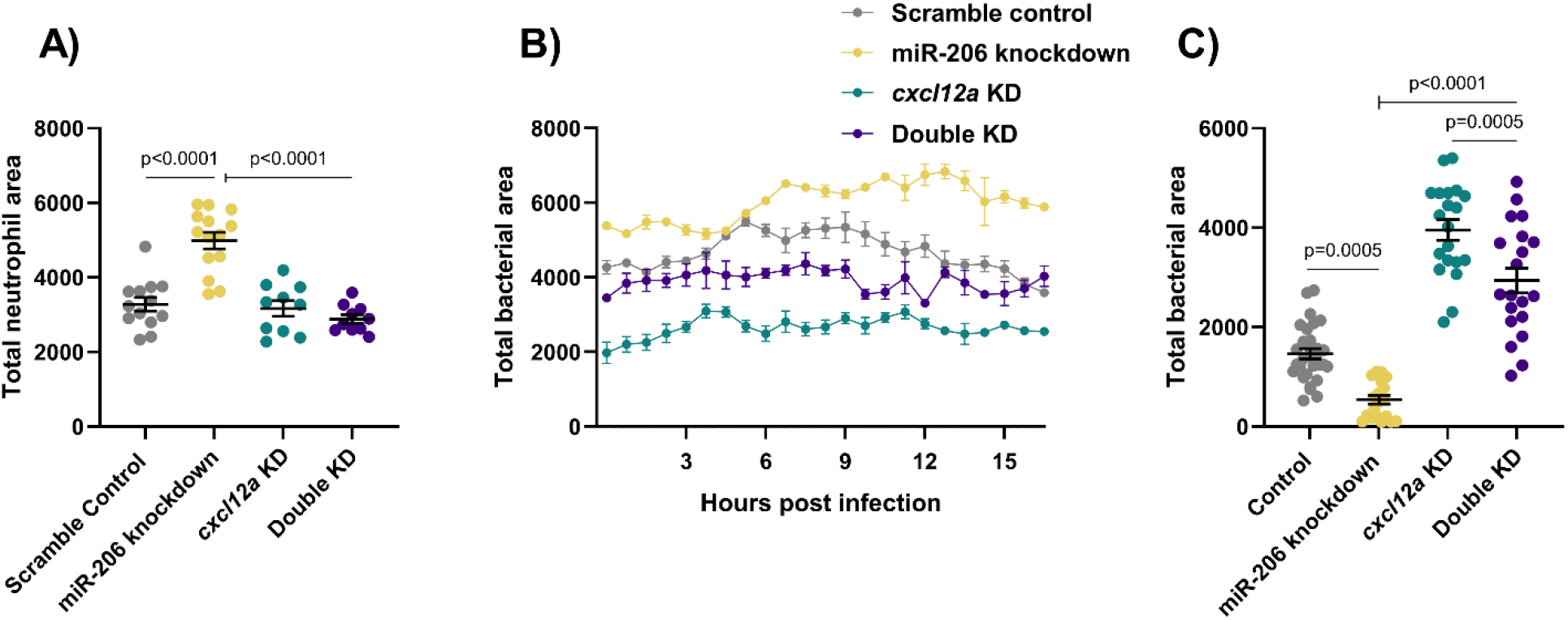
Cxcl12a knockdown places the Cxcl12/Cxcr4 signalling axis downstream of miR-206. (A) Whole body neutrophil counts at 3 dpi of *cxcl12a* and double (*cxcl12a* and miR-206) knockdown embryos. (B) Measurement of neutrophil recruitment to *M. marinum* following trunk injection in double knockdown embryos. (C) Bacterial burden at 3 dpi in *M. marinum*-infected double knockdown embryos. Each data point represents a single measurement, with the mean and SEM shown. For time-lapse imaging, each data point represents the mean of 4 foci of infection from 4 separate embryos. Bacterial burden analysis was performed on 20-30 embryos per treatment. Graphs are representative of 2 biological replicates.

## Discussion

In this study we have demonstrated an *in vivo* link between infection-induced miR-206 expression and the Cxcl12/Cxcr4 signalling axis in the control of mycobacterial infection. Knockdown of miR-206 resulted in a decreased bacterial burden and improved infection outcome. We attribute the reduced bacterial burden to an increased early neutrophil response from increased *cxcr4b* and *cxcl12a* transcript abundance in miR-206 knockdown animals. We show that host miR-206 is increased by pathogenic *M. marinum* to impede the host Cxcl12/Cxcr4 signalling axis, thereby reducing protective early neutrophil recruitment to the site of infection and aiding the creation of a permissive niche for mycobacterial infection.

As this early protective neutrophil response was specific to virulent intracellular mycobacteria, the observed increase in miR-206 was deemed to be ESX1-dependent. We hypothesise that this may be a mycobacteria-driven response to avoid neutrophil phagocytosis and potentially oxidative killing (31). Neutrophils are one of the first immune cells to respond to mycobacterial infection and are capable of both phagocytosing and trapping mycobacteria in neutrophil extracellular traps (32). Therefore, it is not surprising that mycobacteria have evolved a strategy to actively subvert host neutrophil recruitment by reducing Cxcl12/Cxcr4 signalling, limiting the downstream exposure of mycobacteria to phagocytosis and oxidative killing.

Involvement of Cxcr4 and its ligand Cxcl12 in inflammation is well documented (33–35), however, previous studies have largely focused on their role in viral responses. Cxcr4 expression is reduced in the lymphocytes of leprosy patients, but increased in *M. tuberculosis*-infected macrophages (36, 37). Recent work has highlighted Cxcr4 as a mediator of host infection-associated angiogenesis (38), while this study further links the Cxcl12/Cxcr4 signalling axis to virulence-dependent neutrophil recruitment during mycobacterial infections. Cxcr4 has also been shown to participate in pathogen immune evasion via interaction with TLR2, suggesting it may play an active role in other aspects of mycobacterial pathogenesis (39).

Our data provide precedence for cross-species conservation of host miRNA responses to mycobacterial infection. Although current understanding of the role of miR-206 in bacterial infections has been limited to date, previous investigation using *M. tuberculosis* infection of THP-1 cells revealed a similar infection-induced upregulation of miR-206 *in vitro* (11). Our *in vivo* model of mycobacterial infection has allowed the interrogation of neutrophil responses as a downstream cellular response controlled by miR-206.

While we have demonstrated the effect of miR-206 expression on *cxcl12a* and *cxcr4b*, other genes targeted by miR-206 may also contribute to host control of mycobacterial infection. One validated target of miR-206, Vegf, plays a significant role in the later development of granulomas during mycobacterial infection, consistent with the late downregulation of miR-206 that we observed (24, 26, 40). Infection-induced Vegf signalling results in an aberrant angiogenesis programme which favours mycobacterial growth and spread (30, 41-43). This effect may be synergistic with Cxcl12/Cxcr4 signalling, which supports granuloma-associated angiogenesis through a Vegf-independent mechanism (38).

In addition, suppression of *elmo1* by miR-206 may further contribute to the immune avoidance associated with the infection-induced increase in miR-206. Recent investigations have revealed a role for Elmo1 in neutrophil migration and engulfment of apoptotic cells (44) and this has been linked to enhanced intracellular mycobacterial growth (45). Increased transcription of Elmo1 following miR-206 knockdown is likely to increase neutrophil mobility during infection in cooperation with increased Cxcl12/Cxcr4 signalling.

miR-206 may also act on *timp3* to inhibit the activity of Mmp9 during mycobacterial infection (11), preventing macrophage recruitment and granuloma formation in our miR-206 knockdown model (46). However, dissecting the miR-206-Mmp9 interaction may require a different experimental platform to determine if reduced *mmp9* expression is a result of transcriptional feedback from its inhibitor Timp3 or caused by the reduced bacterial burden. This may prove to be an additional pathway modulated by miR-206 during infection, acting to alter disease progression and highlights the complex interaction of miRNA and their multiple targets.

The final potential target gene we profiled, *mmd*, may also be of significance through the positive regulation of macrophage activation and downstream cytokine signalling cascades (47).

In summary, we have identified potential target genes of miR-206 which may be biologically active during mycobacterial infection. We have demonstrated a link between infection-associated upregulation of miR-206 and suppression of neutrophil recruitment to the site of pathogenic mycobacterial infection involving the Cxcl12/Cxcr4 signalling pathway. This host response to infection by pathogenic mycobacteria appears to be conserved across host-pathogen pairings and could inform the development of biomarker or therapeutic strategies.

## Methods

### 1. Zebrafish husbandry

Adult zebrafish were housed at the Centenary Institute and experiments were approved by Sydney Local Health District AWC Approval 17-036. The embryos were obtained by natural spawning and were raised in E3 media and maintained at 28-32°C.

### 2. Zebrafish lines

Zebrafish were AB strain. Transgenic lines used were: *Tg(lyzC:GFP)^nz117^* and *Tg(lyzC:DsRed2)^nz50^* were used for neutrophil imaging experiments (48).

### 3. Embryo microinjection with antagomiR

Embryos were obtained by natural spawning and were injected with either miR-206 antagomiR (-CCACACACUUCCUUACAUUCCA-) or a scramble control (-CAGUACUUUUGUGUAGUACAA-) (GenePharma, China) at 200 pg/embryo at the single cell stage and maintained at 32°C.

### 4. miRNA target prediction

Prediction of target mRNA was performed using TargetScan. dre-miR-206-3p was entered into TargetScanFish 6.2 (http://www.targetscan.org/fish_62/), hsa-miR-206 entered into TargetScan 7.2 (http://www.targetscan.org/vert_72/), and mmu-miR-206 entered into TargetScanMouse 7.2 (http://www.targetscan.org/mmu_72/).

### 5. *M. marinum* culture

*M. marinum* was cultured and quantified as previously described (49). *M. marinum* expressing Wasabi or tdTomato fluorescent protein was used for infections.

### 6. UPEC culture

Uropathogenic *Escherichia coli* (UPEC) carrying the mCherry PGI6 plasmid was cultured in LB supplemented with 50 μg/mL of spectinomycin overnight at 37°C with 200 RPM shaking. Bacteria was then further diluted 1:10 with LB + spectinomycin (50 μg/ml) and incubated for 3 hours at 37°C with 200 RPM shaking. 1 mL of culture was centrifuged (16,000 x *g* for 1 minute), and the pellet washed in PBS. Following another centrifugation, the bacterial pellet was resuspended in 300 μl of PBS + 10% glycerol and aliquoted for storage. Enumeration of bacteria was performed by serial dilution on LB + spectinomycin agar plates and culturing at 37°C overnight. Bacterial concentration was determined by CFU counts.

### 7. UPEC plasmid construction

The plasmid pGI6 was constructed by replacing the open reading frame (ORF) of msfGFP in pGI5 (50) with an E. coli codon-optimised ORF for mCherry. The mCherry ORF was first amplified with the forward primer (GCG CCG CCA TGG GTG AGC AAG GGC GAG GAG GAT) and reverse primer (GGC CCG GGA TCC TTA CTT GTA CAG CTC GTC CAT GCC) from the template pIDJL117 (51). The PCR fragment was cloned at NcoI and BamHI in pGI5, thus replacing msfGFP, and the PCR-generated confirmed by sequencing.

### 8. Bacterial infections

Staged at approximately 1.5 dpf, embryos were dechorionated and anesthetised in tricaine (160 μg/ml). Working solutions of *M. marinum* or UPEC (diluted with 0.5% w/v phenol red dye) were injected into either the caudal vein or trunk to deliver approximately 200 CFU *M. marinum* or 250 CFU UPEC. Embryos were recovered in E3 media + PTU (0.036 g/L) and housed at 28°C.

### 9. Crispr-Cas9 mediated knockdown

Embryos were injected at the 1-2 cell stage with 1 nL of Crispr mixture containing 1 μg/μl Guide (g) RNA (Table 1.), 500 μg/mL Cas9. For double knockdowns with Crispr-Cas9 and antagomiR, mixtures contained 1 μg/μl gRNA, 100 pg/nL antagomiR (miR-206), and 500 μg/mL Cas9. gRNA was synthesised as previously described (52). Embryos were transferred to E3 containing methylene blue and maintained at 32°C.

**Table 1.**
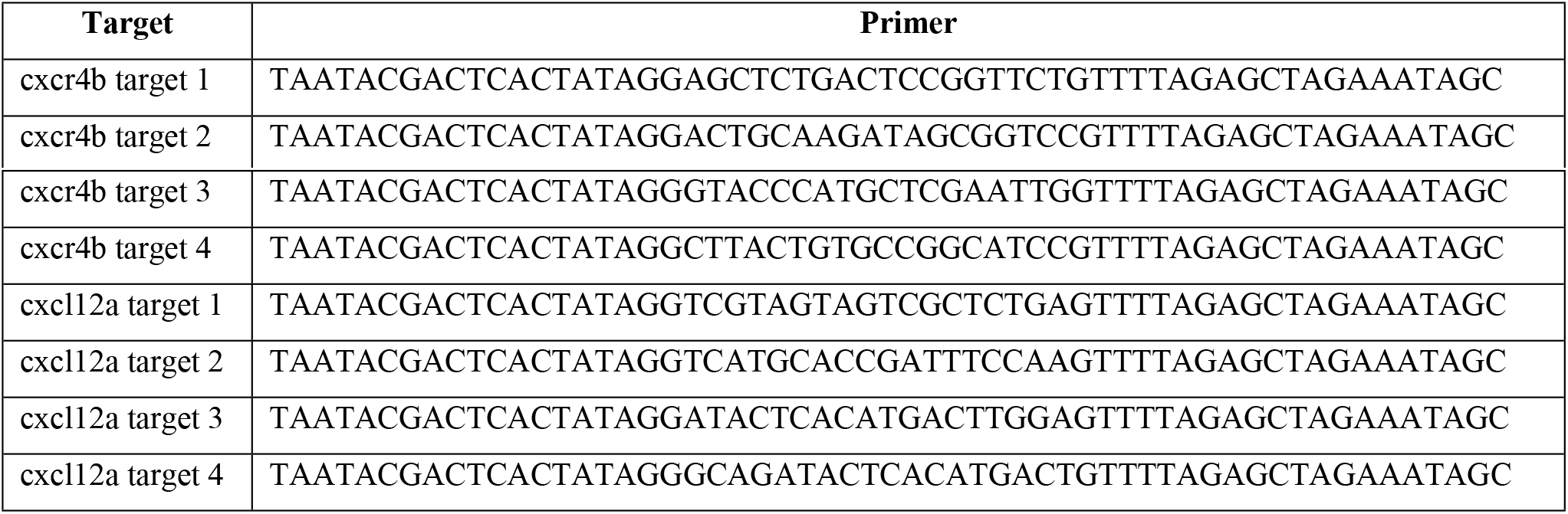
Guide RNA sequences used for Crispr-Cas9 mediated knockdown experiments

### 10. Gene expression analysis

Groups of 10 embryos were lysed and homogenised using a 27-gauge needle in 500 μl Trizol (Invitrogen) and RNA extracted as per the manufacturer’s instructions. cDNA was synthesised from 500 ng RNA using the miScript II RT kit (Qiagen) with HiFlex buffer. qPCR was carried out on an Mx3000p Real-time PCR system using Quantitect SYBR Green PCR Mastermix and primer concentration of 300 nM (Table 2.). For miRNA qPCRs, the miScript Universal Primer was used alongside miR specific miScript primer assays (miR-206 cat. no. MS00001869 and U6 cat. no. MS00033740; Qiagen).

**Table 2.**
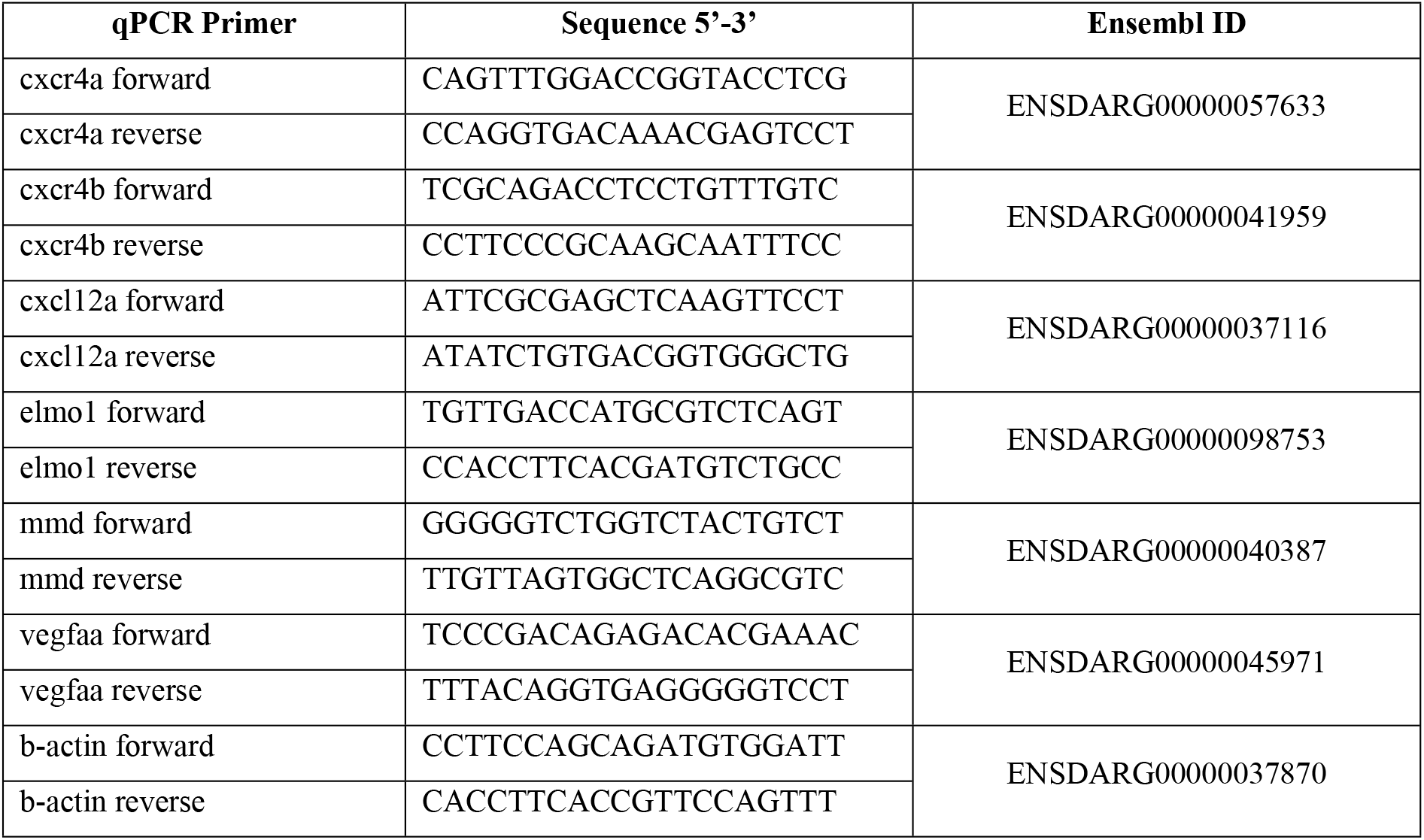
qPCR primer sequences

Cycling conditions for miRNA were: 95°C for 15 minutes; 40 cycles of 95°C for 20 seconds, 56°C for 30 seconds, 72°C for 30 seconds with fluorescence data acquisition occurring at the end of each cycle, followed by 1 cycle of 95°C for 1 minute, 65°C for 30 seconds, and 97°C for 30 seconds. For mRNA, the conditions were: 95°C for 15 minutes; 40 cycles of 94°C for 15 seconds, 55°C for 30 seconds, 70°C for 30 seconds with fluorescence data acquisition occurring at the end of each cycle, followed by 1 cycle of 95°C for 1 minute, 65°C for 30 seconds, and 97°C for 30 seconds.

U6 or β-actin was used as an endogenous control for normalisation and data analysed using the 2^-ΔΔ^ Ct method.

### 11. AMD3100 treatment

Embryos were treated with 20 μM AMD3100 (Sigma-Aldrich), a pharmacological CXCR4 antagonist, dissolved in water and refreshed daily.

### 12. Static imaging and burden analyses

Live imaging was performed on anaesthetised embryos on a depression microscope slide. Images were acquired using a Leica M205FA Fluorescent Stereo Microscope equipped with a Leica DFC365FX monochrome digital camera (Leica Microsystems, Germany). Images were analysed using ImageJ software to quantify the fluorescent pixel count (49).

### 13. Neutrophil tracking analyses

Time-lapse imaging was performed on a Deltavision Elite at 28°C (GE, USA). Following infection with *M. marinum* into the trunk, embryos were mounted in a 96-well black-walled microplate in 1% low-melting point agarose topped up with E3. Images were captured every 60-180 seconds for 16-24 hours. Analysis was performed using ImageJ software. Briefly, every 10-30 images were analysed for the quantity of neutrophils in a 1000 x 500 μm box around infection foci by quantifying the fluorescent pixel count (total neutrophil area) at each time point.

### 14. Statistics

Statistical analysis was performed in GraphPad Prism (v. 9.0.0). All data was analysed by T-test or ANOVA depending on the number of experimental groups, post-hoc analysis performed using Tukey’s multiple comparisons test. For time-lapse data, group comparisons were computed using the Sidak test. Outliers were removed prior to statistical analysis using ROUT, with Q=1%.

## Acknowledgements

Dr Angela Kurz of the BioImaging Facility and Sydney Cytometry at Centenary Institute for technical assistance with imaging, Dr Pradeep Manuneedhi Cholan for assistance with imaging, and Dr Elinor Hortle for assistance with image analysis.

## Author contributions

KW, KDS, KMP, ACP, WJB, SHO conceived the experiments. KW and SHO designed the experiments. IGD and TAB created and provided essential reagents. KW performed the experiments. KW and SHO analysed the data. KW and SHO wrote the paper that was read and approved by all authors.

## Supporting information

**S1 video. Neutrophil migration to infection in control embryos.**

**S2 video. Neutrophil migration to infection in miR-206 knockdown embryos.**

**S3 video. Neutrophil migration to sterile wound site in control embryos.**

**S4 video. Neutrophil migration to sterile wound site in miR-206 knockdown embryos.**

